# Rapid Anthropocene realignment of allometric scaling rules

**DOI:** 10.1101/2020.11.12.379248

**Authors:** Luca Santini, Nick J.B. Isaac

**Author notes:** **Author contributions:** LS conceived the original idea, collected all data, performed the analyses, and drafted the original version of the manuscript. NJBI made substantial contributions to the framing of the study, definition of the methodology, interpretation of the results, and the writing.

## Abstract

The negative relationship between body size and population density (SDR) in mammals is often interpreted as resulting from energetic constraints. In a global change scenario, however, this relationship might be expected to change, given the size-dependent nature of anthropogenic pressures and vulnerability to extinction. Here we test whether the SDR in mammals has changed over the last 50 years. We show that the relationship has shifted down and became shallower, corresponding to a decline in population density of 32-72%, for the largest and smallest mammals, respectively. However, the SDRs become steeper in some groups (e.g. carnivores) and shallower in others (e.g. herbivores). The Anthropocene reorganization of biotic systems is apparent in macroecological relationships that were previously believed to be immutable, reinforcing the notion that biodiversity pattens are contingent upon conditions at the time of investigation. We call for an increased attention on the role of global change on macroecological inferences.

## Introduction

Macroecology seeks to establish relationships that are informative of nature’s underlying mechanisms. The relationship between species body mass and population density, also known as the size-density relationship (SDR), has long been studied to shed light on how the abundance of animals scales with their size (Damuth 1981; Allen *et al*. 2002; Brown *et al*. 2004; White *et al*. 2007). Studies have shown that a clear negative relationship exists (Damuth 1981), which was initially explained in terms of the scaling between body mass and metabolic rate (Kleiber 1932). Damuth (1981, 1987) noted that the scaling coefficients between body mass and metabolic rate (~0.75) and body mass and population density were inverse (~-0.75), suggesting that the energy flux was invariant to body mass, i.e. energy equivalence rule (Brown *et al*. 2004). The energy-equivalence rule implied an energetic tradeoff between physiological and ecological process, resulting in energy in ecosystems being distributed independently of the size of organisms. This theory was later challenged on empirical and theoretical grounds (Carbone *et al*. 2007; McGill 2008; Isaac *et al*. 2011, 2013; Delong & Vasseur 2012; Munn *et al*. 2013; Nilsen *et al*. 2013). The SDR also varies across trophic levels (Silva *et al*. 1997) - with higher trophic levels typically showing lower intercepts and steeper slopes - and taxa (Isaac *et al*. 2011; Pedersen *et al*. 2017b). Competing explanations for the existence of the SDR in endotherms have focused on differential resource availability and accessibility across trophic levels, and energy conversion efficiency (Damuth 1987; Blackburn *et al*. 1993; Silva *et al*. 1997; Carbone & Gittleman 2002; Ernest *et al*. 2003).

An increasing number of studies suggest that what we know about the natural word is contingent to the conditions at the time of investigation (Santini *et al*. 2017). Large-scale emergent properties are assumed to arise from ecosystems tending to an equilibrium, however equilibria are unlikely static and probably continuously shift (McGill 2011). Many species, for example, may be lagged behind environmental changes due to limited dispersal capacities (Svenning & Skov 2004; Araújo & Pearson 2005), or be locally doomed to extinction due habitat loss and fragmentation but still temporarily present due to slow demographic dynamics (i.e. extinction debts; Diamond 1972; Tilman *et al*. 1994). On top of this, the Anthropocene is characterized by a rapid reshuffling of biological communities (Dornelas *et al*. 2014; McGill *et al*. 2015), potentially disrupting the equilibria under which patterns such as the SDR emerged. Examples are the Bergmann rule that has been altered by the extinction of large-bodied species in temperate areas (Faurby & Araújo 2016; Rapacciuolo *et al*. 2017; Santini *et al*. 2017), patterns of geographic range and species richness that appear shaped at least as much by human factors as climate species and biogeography (Di Marco & Santini 2015; Torres Romero & Olalla Tárraga 2015). Similarly, mammal abundance has been shown to increase (Tucker *et al*. 2020) and movements to decrease in anthropogenic areas (Tucker *et al*. 2018), and activity patterns altered globally with nocturnal activity increasing in more disturbed environments (Gaynor *et al*. 2018), and biogeographic realms that change depending on whether introduced and extinct species are considered in the clustering algorithm (Bernardo-Madrid *et al*. 2019). Analogously, authors have claimed that deviations from the energy-equivalence rule may result from historical reassembly of mammal communities (Munn *et al*. 2013).

There are several reasons to expect that the SDR is unstable over time. Abundance can change in response to direct persecution, harvesting (Benítez-López *et al*. 2017), habitat degradation and fragmentation (Pfeifer *et al*. 2017), supplemental resources (Yirga *et al*. 2013), predation or competition release (Terborgh 2015), or human shield effects (Berger 2007). At the same time, these changes are uneven with respect to body mass. Larger species have shown to be intrinsically more at risk of extinction because of diminished resilience, or because are preferentially affected or targeted by human actions (Purvis *et al*. 2000; Cardillo *et al*. 2005). Small and large species also tend to be threatened by different factors, e.g. small species in mammals appear to be more sensitive to habitat loss and degradation, whereas large species to be more sensitive to over-harvesting (Gonzalez-Suarez *et al*. 2013; Ripple *et al*. 2017). Mammal and bird faunas are projected to become increasingly dominated by small-bodied species (Cooke *et al*. 2019).

The action of these anthropogenic pressures, simultaneously and/or heterogeneously, creates potential for both the intercept and slope of the SDR to shift over time. By contrast, theory emphasises fixed physiological and environmental constraints as the primary forces responsible for allometric scaling parameters, so the null expectation would be that these constraints somehow overcome or balance out the effects of anthropogenic forcing (Fig 1a). Anthropogenic pressures have the potential to exert directional pressure on both parameters of the SDR: a general reduction in wild biomass reduce the intercept (Fig 1b), whereas size-dependent declines could lead to a reduction in the slope (Fig. 1c), although these are not mutually exclusive (Fig. 1d). These changes might be further complicated by extinctions (resulting the removal of species from the SDR) and trophic interactions (e.g. reduction in predation pressure and competitive release).

**Fig. 1.**
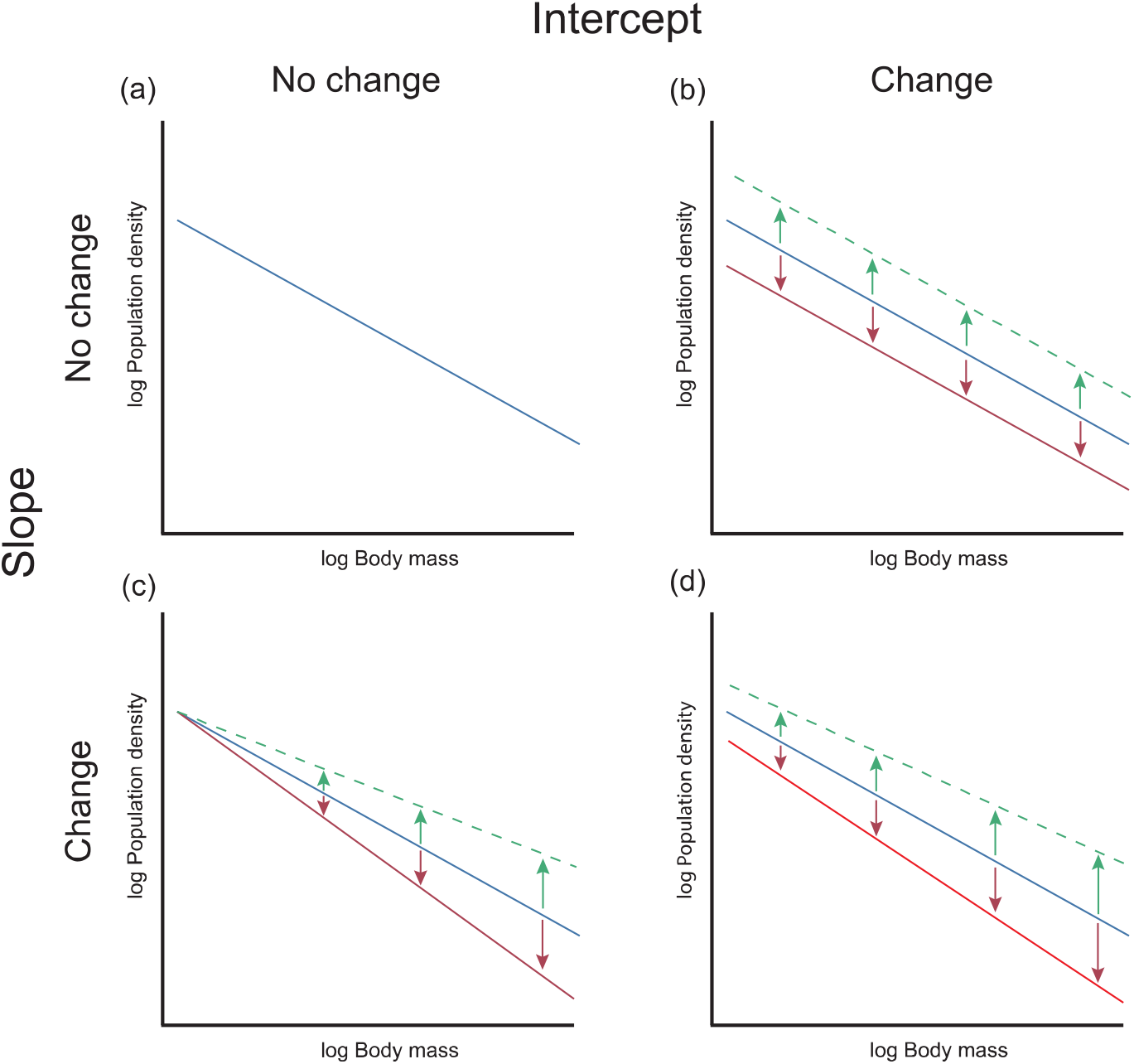
Conceptual representation of the four competing hypotheses on the shift in intercept and slope of the relationship between body mass and population density. (a) Average population densities have remained unchanged and the relationship does not change; (b) Species of different size change in population density at a similar rate, i.e., only intercept change; (c) Only large species change in population density, i.e., only the slope changes; (d) All species change in population density but at different rates depending on their body mass, i.e., the intercept and the slope both change.

Here we assess if and how the SDR has shifted in time across a 50-year period that coincided with unprecedented transformation of natural habitats. We focus on mammals on which SDR investigation has largely focused (Damuth 1981, 1987; Pedersen *et al*. 2017a), and for which population density data have been collected for a long time and body mass data are largely available (Santini *et al*. 2018). Because species of different trophic groups exhibit different relationships (Silva *et al*. 1997; Carbone & Gittleman 2002) and can be threatened by, or benefit, from different environmental changes (Berger 2007; Estes *et al*. 2011; Terborgh 2015; Cooke *et al*. 2019), we also assess if and how these shifts vary across diet categories. Our general hypothesis is that anthropogenic pressures have led to systematic shifts in the parameters of the SDR over the course of a human lifetime, with a general decrease in both intercept and slope parameters. We expect this shift to be particularly evident in carnivores where large species have suffered strong declines (Ripple *et al*. 2014).

## Methods

### Data

We extracted all population density estimates for mammals from an updated unpublished version TetraDENSITY database, totaling 16,738 estimates for 832 species (Santini *et al*. 2018). We only retained data for which the sampling method was reported, and year and location of sampling were known. We excluded all density estimates referred to non-native species. When density estimates were the results of sampling across several years (e.g. mark-recapture studies in large carnivores; 2-4 years), we took the middle year as reference sampling time. This dataset resulted in 13,624 population density estimates for 749 species of mammals. The sampling years spanned between 1918 and 2019, but were mostly distributed between 1970 and 2019 (Fig. 2). In order to avoid possible biases due to limited sampling before 1970, we reduced the dataset to density estimates collected between 1970 and 2020. This reduced the datasets to 13,040 estimates for 736 species in mammals.

**Fig. 2.**
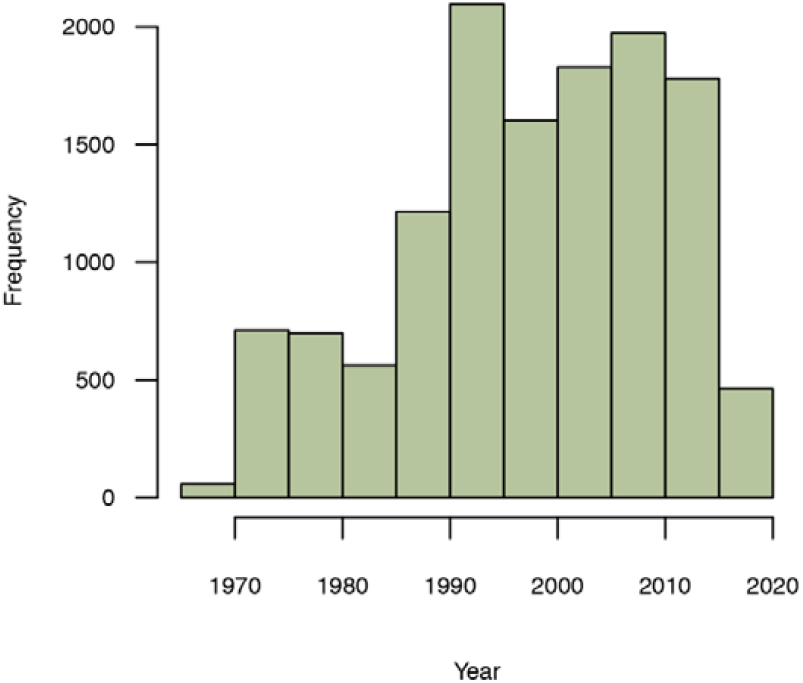
Distribution of the sampling years of the population density estimates mammals (b).

**Fig. 3.**
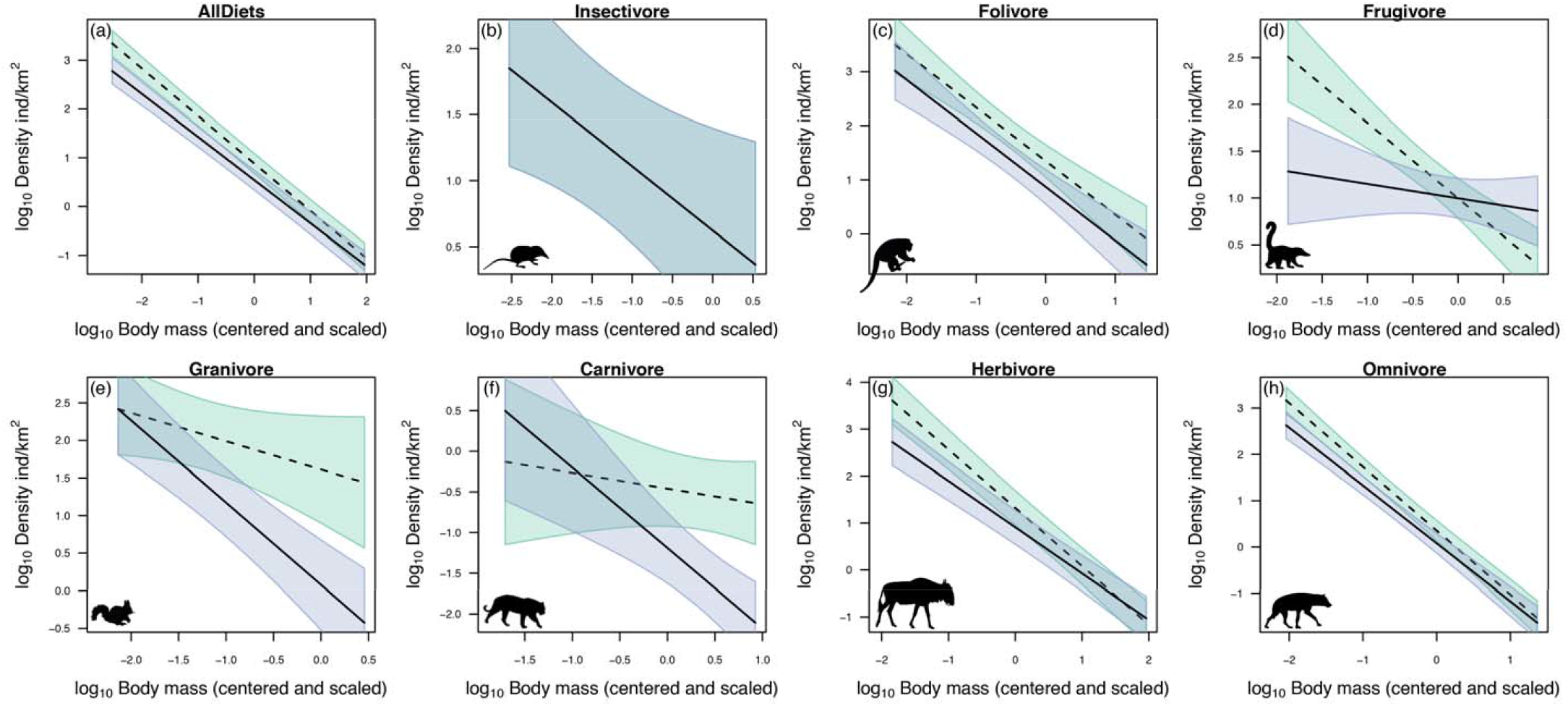
Predicted change in average population density of mammals along the full range of body mass. Dashed line = 1970; Solid line = 2020. (a) all diet categories, (b) insectivores, (c) folivores, (d) frugivores, (e) granivores, (f) carnivores, (g) herbivores, (h) omnivores.

**Fig. 4.**
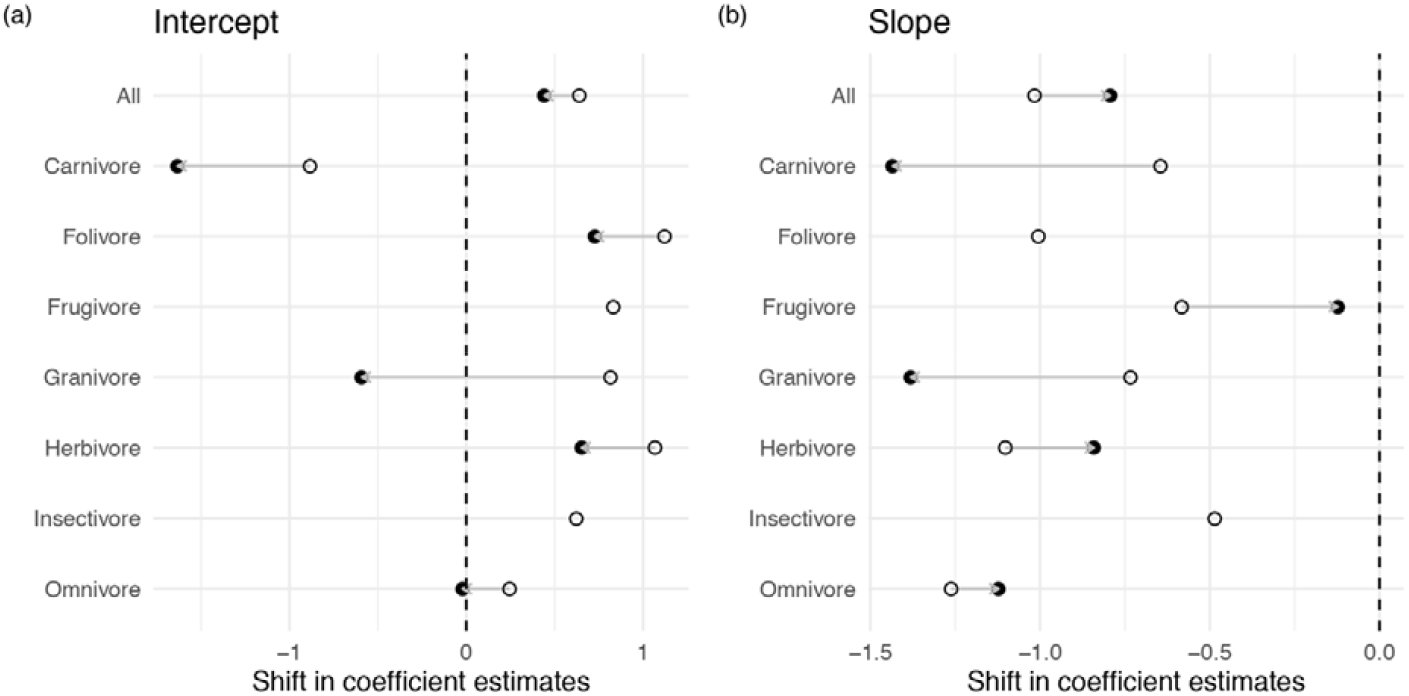
Change in model coefficients for 1970 (empty points) and 2020 (full points) estimated based the additive term of the ‘Year’ variable (change in intercept) and the interaction term between ‘Body mass’ and ‘Year’ (change in slope).

Body mass and diet information for all mammals were extracted from EltonTraits database v. 1.0 (Wilman *et al*. 2014). EltonTraits provides proportions of diet items in mammals, which we used to classified mammals into three seven diet categories as in Silva et al. (1997). Species with >=90% vertebrate-based diet were classified as carnivores (n=770, sps=34) and those >=70% invertebrate-based diet as insectivores (n=300, sps=59). Species with <20% of animal-based diet were classified as frugivore if with >=40% fruit in diet (n=1571, sps=159) and as granivores if with >=30% seed-based diet (n=785, sps=63). Although EltonTraits does not differentiate between leaves and grass, we further distinguished herbivores from folivores by considering herbivores species with >=80% plant-based diet belonging to cetartiodactyla and perissodactyla (n=6119, sps=184), and as folivores species not belonging to these groups (with the exception of Giraffidae) with >=60% plant-based diet (n=545, sps=53). All species that did not fit the previous two categories were classified as omnivores (n=2950, sps=184).

To account for phylogenetic relatedness in the modelling, we used the phylogeny in Upham et al. (2019). We extracted 1000 random trees and generated a majority-rule consensus setting the frequency of in the sample with which each clade or bipartition is encountered at 0.8. We then computed a principal component analysis on the phylogenetic distance matrix of the consensus tree, and extracted the first 5 eigenvectors explaining >99% of the total variance.

All data used in this study are made available as part of the supporting information.

### Modelling approach

We fitted a set of generalized mixed effect linear models (GLMM) using a gaussian distribution to test and compare our hypotheses (Table 1). All models included a random effect at the level of species to control for the effect of pseudo-replicas and possible species turnover through time. We also included a random effect to control for different sampling methodologies broadly classified into eight categories: censuses (‘complete’ counts, which assume full detection of individuals), distance sampling (including different algorithms and sampling design), home range extrapolation (derived from home range area estimation), mark–recapture (including different algorithms and capture approaches), N-Mixture models, Random Encounter models, incomplete counts (any incomplete count that is extrapolated to a larger area), trapping (removal methods, indicate the minimum number known to be alive).

**Table 1.** Hypotheses tested and corresponding fixed effect model formulas. PD = log_10_ Population density; BM = log_10_ Body mass; Year = Sampling year.

| Hypothesis | Model | Formula | Explanation |
| --- | --- | --- | --- |
| Null hypothesis | Null | PD ~ BM | Population densities are constant over time |
| Intercept shifts | IntOnly | PD ~ BM + Year | Population densities shift over time |
| Slope shifts | SlopeOnly | PD ~ BM + Year:BM | Size-dependent shift in population density where the intercept does not change |
| Intercept and slope shift | IntSlope | PD ~ BM + Year + Year:BM | Population densities shift over time but change is size-dependent |

The model representing our null hypothesis (Fig. 1a) only included body mass (Table 1). To assess whether the intercept has changed, we tested the additive effect of time by including sampling year as a fixed effect. To assess whether the slope has changed, we tested the effect of time along the body mass range by included an interaction term between sampling year and body mass (Table 1). In order to test whether our hypotheses were only valid for a specific diet category, we repeated the model selection for each diet category separately.

Models were compared using the Akaike Information Criterion (AIC). We also present results using the Bayesian Information Criterion (BIC), which tends to be more conservative than AIC and is thus considered more suitable for very large sample sizes (Raffalovich *et al*. 2008). We tested the residuals of the selected model for spatial autocorrelation using Moran I and phylogenetic autocorrelation using Pagel’s lambda and 100 random phylogenetic trees from Upham et al. (2019). We also plot the residuals against time to assess a possible temporal autocorrelation effect in the residuals.

## Results

The most supported model explaining the SDR across all species was the one accounting for a shift in both intercept and slope (IntSlope model; Table 2, S1). The same model was the most supported according to AIC in all diet groups except for insectivores and folivores (Table 2, S1). While in both insectivores and folivores the IntSlope model was also competitive (within 2-AIC units from the best models), in these cases we selected the most parsimonious models, the Null and the Intercept-only model, respectively. (Table 2, S1).

**Table 2.** Comparison of models tested through AIC and BIC. n = sample size; df = number of parameters; LL = log-likelihood; Δ = delta AIC or BIC; ω = AIC or BIC weights. The best model according to AIC is highlighted in bold.

| | Model | n | df | LL | AIC | ΔAIC | AICω | BIC | ΔBIC | BICω |
| --- | --- | --- | --- | --- | --- | --- | --- | --- | --- | --- |
| AllDiets | **IntSlope** | **13028** | **7** | **-12512.21** | **25038.42** | **0** | **0.99** | **25090.74** | **0** | **0.63** |
|  | IntOnly |  | 6 | -12517.49 | 25046.99 | 8.57 | 0.01 | 25091.83 | 1.09 | 0.37 |
|  | SlopeOnly |  | 6 | -12572.69 | 25157.39 | 118.97 | 0 | 25202.24 | 111.5 | 0 |
|  | Null |  | 5 | -12579.61 | 25169.22 | 130.8 | 0 | 25206.59 | 115.85 | 0 |
| Insectivore | IntSlope | 300 | 7 | -345.49 | 704.99 | 0 | 0.5 | 730.91 | 6 | 0.04 |
|  | **Null** |  | **5** | **-348.2** | **706.39** | **1.41** | **0.24** | **724.91** | **0** | **0.82** |
|  | SlopeOnly |  | 6 | -347.57 | 707.14 | 2.16 | 0.17 | 729.37 | 4.45 | 0.09 |
|  | IntOnly |  | 6 | -348.17 | 708.35 | 3.36 | 0.09 | 730.57 | 5.66 | 0.05 |
| Folivore | **IntOnly** | **545** | **6** | **-488.38** | **988.77** | **0** | **0.7** | **1014.57** | **0** | **0.83** |
|  | IntSlope |  | 7 | -488.35 | 990.71 | 1.94 | 0.27 | 1020.81 | 6.24 | 0.04 |
|  | SlopeOnly |  | 6 | -492.17 | 996.34 | 7.57 | 0.02 | 1022.14 | 7.57 | 0.02 |
|  | Null |  | 5 | -493.54 | 997.07 | 8.3 | 0.01 | 1018.58 | 4 | 0.11 |
| Frugivore | IntSlope | 1571 | 7 | -1382.51 | 2779.02 | 0 | 0.6 | 2816.54 | 4.04 | 0.07 |
|  | **SlopeOnly** |  | **6** | **-1384.17** | **2780.35** | **1.32** | **0.31** | **2812.5** | **0** | **0.51** |
|  | IntOnly |  | 6 | -1385.57 | 2783.14 | 4.12 | 0.08 | 2815.3 | 2.79 | 0.13 |
|  | Null |  | 5 | -1388.39 | 2786.79 | 7.76 | 0.01 | 2813.58 | 1.08 | 0.3 |
| Granivore | **IntSlope** | **785** | **7** | **-615.27** | **1244.55** | **0** | **0.99** | **1277.2** | **0** | **0.88** |
|  | IntOnly |  | 6 | -620.74 | 1253.49 | 8.94 | 0.01 | 1281.48 | 4.28 | 0.1 |
|  | SlopeOnly |  | 6 | -623.4 | 1258.8 | 14.26 | 0 | 1286.8 | 9.59 | 0.01 |
|  | Null |  | 5 | -627.36 | 1264.73 | 20.18 | 0 | 1288.05 | 10.85 | 0 |
| Carnivore | **IntSlope** | **770** | **7** | **-642.52** | **1299.05** | **0** | **0.82** | **1331.58** | **1.42** | **0.3** |
|  | IntOnly |  | 6 | -645.14 | 1302.28 | 3.23 | 0.16 | 1330.16 | 0 | 0.62 |
|  | SlopeOnly |  | 6 | -647.14 | 1306.29 | 7.24 | 0.02 | 1334.17 | 4.01 | 0.08 |
|  | Null |  | 5 | -674.2 | 1358.41 | 59.35 | 0 | 1381.64 | 51.48 | 0 |
| Herbivore | **IntSlope** | **6119** | **7** | **-6069.78** | **12153.57** | **0** | **1** | **12200.6** | **0** | **0.95** |
|  | IntOnly |  | 6 | -6077.04 | 12166.09 | 12.52 | 0 | 12206.4 | 5.8 | 0.05 |
|  | Null |  | 5 | -6090.46 | 12190.93 | 37.36 | 0 | 12224.52 | 23.92 | 0 |
|  | SlopeOnly |  | 6 | -6090.27 | 12192.54 | 38.97 | 0 | 12232.85 | 32.25 | 0 |
| Omnivore | **IntSlope** | **2938** | **7** | **-2608.6** | **5231.2** | **0** | **0.88** | **5273.1** | **1.91** | **0.28** |
|  | IntOnly |  | 6 | -2611.64 | 5235.27 | 4.08 | 0.12 | 5271.19 | 0 | 0.72 |
|  | SlopeOnly |  | 6 | -2616.86 | 5245.73 | 14.53 | 0 | 5281.64 | 10.45 | 0 |
|  | Null |  | 5 | -2630.33 | 5270.66 | 39.46 | 0 | 5300.59 | 29.4 | 0 |

Projecting the model for the 1970 and for 2020, the SDR across all diet categories shifted down and became shallower. However, the qualitative nature and extent of the change varied across diet categories. In herbivores, frugivores and omnivores, the SDR also shifted down and became shallower, whereas the SDR in granivores and carnivores shifted down and became steeper. In folivores the relationship only shifted down, whereas the most parsimonious model for insectivores did not detect any change.

These changes equate to substantial relative changes in the average abundance of species when transformed back to natural scale. Considering the extreme body masses per dataset, on average small species decreased by 72% whereas large by 32%. In folivores density decreased homogeneously by 67%. In frugivores density decreased by 94% in small species and increased by 273% in large ones. Similarly, small herbivores decreased by 87% and large increased by 24%. The density of small granivores remained similar, but the density of large species decreased by 98%. Small carnivores increased by 323% and large carnivores decreased by 96%. Finally, omnivores decreased by 71% and 20% for small and large species, respectively.

BIC-based selection concurs for all groups except in frugivores, carnivores and omnivores. In frugivores the null model is more supported by BIC, whereas in carnivores and omnivores the most supported models by AIC are also ranked as competitive by BIC, but the intercept-only model would be preferred as more parsimonious.

We did not detect an effect of phylogenetic, spatial or temporal autocorrelation in the residuals of the selected models (Table S2, Fig. S1-S2).

## Discussion

Our results suggest that the relationship between body mass and population density, which has long been subject of macroecological investigation (Damuth 1981; Allen *et al*. 2002; Brown *et al*. 2004; White *et al*. 2007; Isaac *et al*. 2013), has changed within a relatively short time frame with mammals declining in density on average. This change is supported in several independent groups of species with different diets. These findings demonstrate that the Anthropocene reorganization of biotic systems is apparent in macroecological relationships that were previously believed to be immutable, casting doubts on our ability to identify “natural” patterns reflecting pure ecological mechanisms.

Overall our results suggest that the macroecological investigation of SDR has been somewhat biased by current biodiversity trends. For example, diet categories exhibit different intercepts that have been interpreted as a matter of resource availability (Silva *et al*. 1997), yet the extent of these difference changes over time, therefore possibly reflecting additional processes (e.g. human exploitation, habitat degradation, resource depletion or supplementation, or human-wildlife conflicts). The somewhat unexpected result of a larger decrease in small than large herbivores may mirror the larger decrease in large carnivores, indicating an overall predation release effect (Hoeks *et al*. 2020). Marquet (2002) noted that the steeper SDR relationship in carnivores has long puzzled ecologists. While this has been explained considering that large preys tend to be distributed less evenly therefore resulting less accessible than small ones (Carbone *et al*. 2007), our results suggest that the observed steeper slope may be the result of the widespread decline of large carnivores (Ripple *et al*. 2014) - and possibly also influenced by the increase of mesocarnivores (Prugh *et al*. 2009; Ritchie & Johnson 2009) - a process that presumably started long before 1970. In fact, while the change in 50 years only is striking, it is likely that changes in species average abundance started much earlier than the first macroecological investigations of the body mass - density relationship (Damuth 1981). We can only speculate that pre-Anthropocene” regression lines are probably higher and shallower than observed in available empirical data. In principle, if we were able to estimate the decline with actual anthropogenic pressures, it would be possible to estimate the “natural” regression line (Santini *et al*. 2017). Yet, this presents several complications in this case, especially considering that every species responds differently to human pressures. Besides, species abundances in ecosystems are strictly linked, so species may increase or decrease in abundance because of the relative abundance of other coexisting species (Terborgh 2015). Other authors have argued that natural baseline could be estimated by focusing on historically less impacted regions of the world (e.g. Africa; Floejgaard *et al*. 2020). A different and promising avenue of research, would be estimating “natural” abundances through the use of mechanistic ecosystem models, e.g. the Madingley model (Harfoot *et al*. 2014). Interestingly, validation of the emergent properties of this model exhibited higher densities than observed in reality, which has been attributed to a number of factors, among which the overall decrease in natural population densities due to human impact (Harfoot *et al*. 2014). Further improvement of global ecosystem models may open up possibilities to estimate natural baselines for macroecology.

The mismatch between AIC-based and BIC-based selection reflects the highly noisy nature of population density data. While our sample sizes were undoubtedly large, detecting a temporal trend within 50 years in a dataset including many species and their trends require a considerable sample size, so uncertainty remain regarding the change in SDR in frugivores, omnivores and carnivores. The likelihood of their models including an interactive term was higher, but not sufficiently high to undeniably justify increased model complexity.

It is possible that our dataset of empirical density estimates suffers of temporal biases which may partly explain our results. For example, it was noted that global mass – density relationship (including species from different ecosystems) may be biased toward higher densities, as researchers often study populations where they are mostly abundant (Marquet *et al*. 1995). This phenomenon could change in intensity over time, perhaps because researchers may have become better able to estimate low-density populations. Similarly, species sampled over time may have slightly changed, for example shifting research attention from management of abundant species to detection of rare species requiring conservation attention. The species-level random effects controls effects due to species turnover, but would not capture a shift in focus from high-density to low-density populations within the geographic range of low-density species. Yet, the consistent results across different diet categories and using different methods suggests this possible effect cannot be the only driver of changes in the SDR parameters over time. While we cannot provide a conclusive answer on the causes of these changes, they ultimately indicate that macroecologists should be very careful in drawing conclusions on macroecological patterns ignoring that these may have been severely modified in recent time, and may be still in the process of changing. Macroecology may thus be unable to get better estimates of pre-Anthropocene allometric parameters just by collecting more data. Perhaps, pre-Anthropocene parameters should not be considered as a natural baseline either, as there is no evidence emergent properties were more stable before humans.

Our understanding of natural world is biased to a compromised situation. The collection of large databases including data collected over long time spans may help us to capture these biases and possibly correct for them. It is crucial that the effect of humans is increasingly considered while assessing and interpreting natural patterns and their causes.

